# Sex Differences in Middle Cerebral Artery Reactivity and Hemodynamics Independent from Changes in Systemic Arterial Stiffness in Adult Sprague-Dawley Rats

**DOI:** 10.1101/2024.11.25.625290

**Authors:** Jonathan W. Ray, Xuming Sun, Nildris Cruz Diaz, Victor M. Pulgar, Liliya M. Yamaleyeva

**Author notes:** Denotes co-corresponding authors Correspondence: Liliya M. Yamaleyeva, Victor M. Pulgar.

## Abstract

Young women are protected against cerebrovascular disease compared with men. However, the underlying mechanisms of sex differences in cerebrovascular function are not well understood. In this study, we determined whether sex differences in middle cerebral artery (MCA) reactivity are accompanied with changes in cerebral or systemic arterial resistance and stiffness in adult 25-week-old Sprague-Dawley (SD) rats. No differences in systolic or diastolic blood pressures were observed between sexes. Heart rate was higher in the female versus male SD. Left MCA pulsatility index (PI) was lower in female versus male SD, while no differences in left intracranial internal carotid artery (ICA) PI was observed between sexes. There were no differences in thoracic aorta or left common carotid artery pulse wave velocity (PWV) between sexes. In isolated MCA segments, female left MCA had lower contraction to potassium, but similar maximal contraction and sensitivity to thromboxane A2 receptor agonist U46619. Pre-incubation with indomethacin lowered maximal response and sensitivity to U46619 in male MCA but not female MCA suggesting that vasoconstrictor prostaglandins may have a greater role in male MCA vs. female MCA. Endothelial nitric oxide synthase (eNOS) and vascular smooth muscle layer thromboxane A2 receptor immunoreactivity were greater in female versus male SD. We conclude that sex differences in the MCA reactivity are associated with a differential functional profile of MCA in adult SD rats independent from changes in systemic PWV.

**NEW & NOTEWORTHY:** Despite the debilitating effects of cerebrovascular disease in women, the basis for sex differences in cerebrovascular dysfunction remains incompletely understood. Our study demonstrated that middle cerebral artery reactivity and hemodynamics are not accompanied by changes in central pulse wave velocity in mature adult Sprague-Dawley rats suggesting different mechanisms underlying baseline vascular reactivity of cerebral versus systemic arterial beds.

## INTRODUCTION

Cerebrovascular disease is a major risk factor for the development of cognitive impairment. It is now well understood that women are at greater risk for cerebrovascular disease (1). Women have a higher lifetime risk of stroke, a higher risk for Alzheimer’s disease, and greater cognitive ability deficits in older age compared to men (2-5). Despite the debilitating effects of cerebrovascular disease in women, the basis for sex differences in cerebrovascular dysfunction remains incompletely understood. Therefore, it is imperative to further our understanding of the vascular changes contributing to increased risk of cerebrovascular impairment in women to identify novel diagnostic and therapeutic targets.

The middle cerebral artery (MCA) is a major cerebral artery and the largest branch of the internal carotid artery (6). It is one of the most common pathologically affected cerebral blood vessels and therefore remains a focal point of research on cerebrovascular disease (6). Sex differences in structure and function of the MCA are reported and likely contribute to the sex differences surrounding cerebrovascular disease severity and onset. Previous works shows that female MCAs exhibit smaller internal diameter, less vascular smooth muscle cells (VSMCs), and increased collagen and elastin content as well as impaired cerebral blood flow autoregulation (7). Additionally, greater peripheral arterial stiffness in women has been linked to increased pulsatile cerebral blood flow, a known contributor to the pathogenesis of cerebrovascular disease (8-17). Young women are relatively protected from cerebrovascular disease compared with older women or men, however there is a rapid doubling of cerebrovascular disease in women in the decade following menopause (1, 18-20). The milieu of vasoactive sex hormones may contribute to sex differences in MCA dysfunction (1, 18-20). Indeed, sex hormones (including estrogens, progestins, and androgens) have been shown to exert influence over numerous functions of cerebral blood vessels including maintenance of cerebrovascular tone and cerebral blood flow, angiogenesis, vascular remodeling, inflammation, and maintenance of the blood brain barrier (19). Furthermore, oral contraceptive use and hormone replacement therapy are sex-specific risk factors for stroke in women (1, 21, 22).

Transcranial Doppler (TCD) ultrasound analysis of the MCA is a non-invasive technique which provides valuable insight into cerebrovascular function in human subjects and laboratory animals (23-25). Pulsatility index (PI), a measure of vascular resistance, is one of the parameters determined by the TCD ultrasound technique: increased PI indicates increased vascular resistance and downstream hypoperfusion (26). MCA PI has been used to predict vascular cognitive impairment in hypertensive patients (27). Since the relationship between central pulse wave velocity and middle cerebral artery pulsatility and reactivity may dependent on sex (28), in this study, we determined sex differences in isolated MCA reactivity and resistance, and systemic arterial stiffness in 25-week-old Sprague-Dawley rats.

## MATERIALS AND METHODS

### Animals

The study was approved by the Institutional Animal Care and Use Committee of the Wake Forest University School of Medicine (A21-057). 25-week-old male and female Sprague-Dawley (SD) rats purchased from Charles River Laboratories (Wilmington, MA, USA) were housed at a constant room temperature, humidity, and light cycle (12:12-h light-dark), fed a standard rodent chow (Lab Diet 5P00 – Prolab RMH 3000, PMI Nutrition International, INC, Brentwood, MO) and given water ad libitum throughout the experimental protocols.

### Blood pressure and heart rate measurements

Systolic (SBP) and diastolic (DBP) blood pressures and heart rate were recorded in trained, conscious rats by the tail-cuff method using the Non-Invasive Blood Pressure (NIBP) Monitor System (Columbus Instruments, Columbus, OH, USA). Data were averaged for each animal and reported as mean ± SEM.

### Transcranial Doppler ultrasound

High-frequency ultrasound was used to determine the pulsatility index (PI) of the left MCA (LMCA) and intracranial portion of the left internal carotid artery (LICA). Animals were placed on a temperature-controlled platform. Temporal hair was removed using a depilatory cream (Nair, Church & Dwight Co., Ewing, NJ). Ultrasound was performed using a Vevo LAZR Ultrasound and Photoacoustic System and LZ250 transducer (FujiFilm, VisualSonics, Toronto, Canada) under 1.5% isoflurane anesthesia. The LMCA is visualized in color Doppler mode by directing the transducer through the rat temporal foramen as previously described (25). Maximum (V_max_), minimum (V_min_), and mean (V_mean_) blood flow velocities were determined by pulse wave Doppler mode and were averaged over three cardiac cycles (**Figure 1A, B**). PI was calculated as follows: PI = V_max_-V_min_/V_mean_. Data were analyzed using Vevo LAB software version 5.7.1 for Windows (FujiFilm VisualSonics, Toronto, Canada).

**Figure 1.**
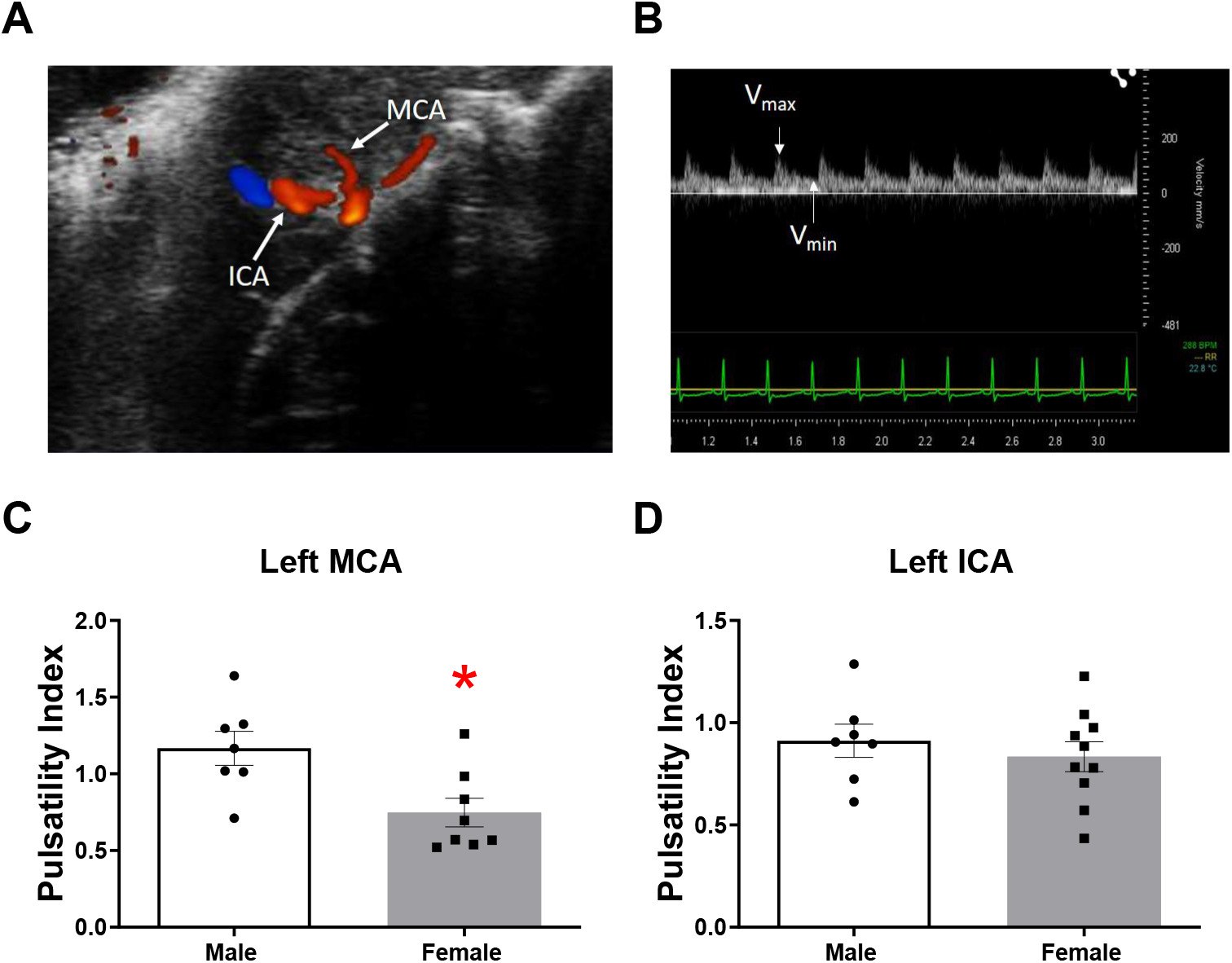
Transcranial Doppler ultrasound and pulsatility index (PI) of left middle cerebral artery (MCA) and left internal carotid artery in adult Sprague-Dawley (SD) rats. Color Doppler ultrasound was used to identify the LMCA which appeared as the first large branch arising from the intracranial internal carotid artery (**Panel A**). Waveforms obtained from pulse wave Doppler analysis provide maximum, minimum, and mean flow velocities which are used to calculate PI (**Panel B**). Left MCA (**Panel C**) and ICA (**Panel D**) PI in 25-week-old male versus female SD rats. Data are mean ± SEM; *p<0.05 vs. male; n=7-10.

### Pulse wave velocity measurements

Pulse wave velocity (PWV) of the thoracic aorta and left common carotid artery (LCCA) were determined as a measure of systemic vascular stiffness (29). Time from R of QRS complex to pulse wave Doppler impulse at the foot of proximal (t_1_) and distal points (t_2_), and the distance between t_1_ and t_2_ are used to calculate the PWV as follows: PWV = d/(t_2_-t_1_) as described by us (30).

### Immunohistochemistry

LMCAs were fixed in 10% formalin and 70% ethanol, embedded in paraffin, and cut into 5-µm sections. Immunostaining was performed using the avidin biotin complex (ABC) method with a diaminobenzene solution used as the chromogen. Antigen retrieval treatment with IHC-TEK Epitope Retrieval Solution (IHC World, Woodstock, MD, USA) was applied at 95-98°C for 40 min. Non-specific binding was blocked in a buffer containing 10% normal goat serum, 0.1% bovine serum albumin, and 1% Triton X-100 in PBS for 30 min. LMCA sections were incubated with rabbit polyclonal cyclooxygenase 2 antibody (COX-2; dilution 1:200; Cayman Chemical, Ann Arbor, MI, USA), mouse monoclonal endothelial nitric oxide synthase antibody (eNOS; dilution 1:200; BD Biosciences, Franklin Lakes, NJ, USA), rabbit polyclonal thromboxane A2 receptor antibody (TxA_2_R; dilution 1:1000; Alomone Labs, Jerusalem, Israel), or rabbit polyclonal prostaglandin I synthase antibody (PGIS; 1:200 dilution; Cayman Chemical, Ann Arbor, MI, USA) and secondary biotinylated goat anti-rabbit or anti-mouse antibodies (dilution 1:400; Vector Laboratories, Newark, CA, USA). Representative images were acquired from each slide with a Mantra Microscope at 40x magnification using Mantra Snap acquisition software (Perkin Elmer, Waltham, MA, USA). Regions of interest (ROI) were defined using the open-source Fiji software (ImageJ, National Institutes of Health). COX-2, TxA_2_R, and PGIS were analyzed in the endothelial and tunica media layers of MCA, while eNOS was analyzed in the endothelial layer of MCA. Intensity of the staining in five ROIs per segment was quantified as described previously by us following the reciprocal intensity method (31-33).

### Vascular reactivity

LMCAs were dissected and mounted between an isometric force transducer (Kistler Morce DSC 6, Seattle, WA, USA) and a displacement device on a wire myograph (Multi Myograph, Model 620M Danish Myo Technologies, Aarhus, Denmark) using two stainless steel wires (diameter 40 μm), using techniques previously described (31, 32, 34). The myograph organ bath (5 ml) was filled with KHB maintained at 37 °C and aerated with 95% O_2_/5% CO_2_. The vessels were washed and incubated for 30 min before the normalization procedure was performed. Arterial segments were normalized to 0.9·L100, with L100 being the internal circumference the vessels would have if they were exposed to a transmural pressure of 100 mmHg. Each arterial segment was stretched in 50 μm steps, internal circumference (L) and wall tension at each stretch level were recorded to produce a resting wall tension-internal circumference curve using the DMT Normalization Module (ADInstruments) (31, 32, 34). Optimal diameters (OD) were calculated as OD=0.9·L100/π. Responses to agonists were recorded after an equilibration period of 30 min. *Response to acetylcholine*. LMCAs were washed and stimulated with a sub-maximal dose of U-46619 (10^−6.5^ M, Cayman Chemical, Ann Arbor, MI, USA), once the contraction was stable a dose response curve to acetylcholine (10^−^10-10^−4^ M) was performed. *Response to the thromboxane analog U-46619*. Contractile response to U-46619 was tested on basal tone. LMCAs were exposed to 13 increasing concentrations of U-46619 (10^−10^-10^−5^ M) applied in half long steps.

### Statistical analysis

The differences between male and female groups were compared using unpaired t-test. The immunostaining for various proteins (**Figure 4**) was analyzed using two-way analysis of variance (ANOVA) followed by the Tukey post-hoc tests (GraphPad Software Inc., La Jolla, CA). All data were presented as mean ± SEM. Data analysis for vascular experiments was performed using GraphPad. Individual experimental data from concentration-response curves for ACh and U-46619 were fitted to the following logistic curve to determine the maximal response and sensitivity Y=bottom + (top-bottom)/ (1+10(LogEC_50_-X)*Hill Slope) where X is the logarithm of the concentration and Y is the response. Basal resting tone and active tone (plus 75mM K+) were expressed as arterial wall tension (AWT) (AWT=force/ 2×length of vessel). Response to ACh was expressed as % of pre-constriction and response to U-46619 was expressed as % of K_MAX_ (maximal response to KCl 75 mM). Sensitivity was expressed as pD2 (pD_2_=−log [EC_50_]). Data are expressed as mean ± SEM. Values of maximum response and sensitivity were compared by Student’s *t*-test. For all experiments, a p-value less than 0.05 was considered statistically significantly different.

## RESULTS

### Physiological Characteristics of 25-Week-Old Male versus Female SD Rats

**Table 1** shows that male SD rats had greater body weight compared with female SD rats. Females had lower heart and mean kidney weight (normalized to tibia length). There was no difference in systolic or diastolic blood pressures between sexes. Males had a greater pulse pressure versus females. However, female SD rats had a greater heart rate versus male SD.

**Table 1.** Physiological Characteristics of 25-Week-Old Male versus Female SD Rats.

|  | Male | Female |
| --- | --- | --- |
| Body Weight, kg | 619.9 ± 37.5 | 349.0 ± 13.7* |
| Brain Weight/Tibia Length, g/cm | 0.4679 ± 0.0164 | 0.4968 ± 0.0075 |
| Heart Weight/Tibia Length, g/cm | 0.3080 ± 0.0133 | 0.2394 ± 0.0102* |
| Mean Kidney Weight/Tibia Length, g/cm | 0.3736 ± 0.0159 | 0.2708 ± 0.0061* |
| Systolic BP (mmHg) | 136.2 ± 4.3 | 128.6 ± 2.5 |
| Diastolic BP (mmHg) | 94.2 ± 3.6 | 94.7 ± 2.3 |
| Pulse Pressure (mmHg) | 42 ± 1.3 | 33.9 ± 1.2* |
| Heart Rate (bpm) | 343.1 ± 10.0 | 438.2 ± 9.1* |

Data are mean ± SEM, \*p<0.05 vs. Male, n=7-11. Bpm, beats per minute; kg, kilogram; g, gram; cm, centimeter. Comparisons made vs. Male by unpaired Student's t test.

### Sex Differences in Left MCA Pulsatility Index in Adult SD Rats

**Figure 1A and B** show Color Doppler signal for left ICA and MCA with a typical wave form of MCA. **Figure 1C** demonstrates that females had lower LMCA PI compared to males. There was no difference in LICA PI between female and male SD (**Figure 1D**). Additionally, there was no difference in blood flow velocity measurements (V_max_, V_min_, and V_mean_) between sexes for either LMCA or LICA (**Figure 2A and B**).

**Figure 2.**
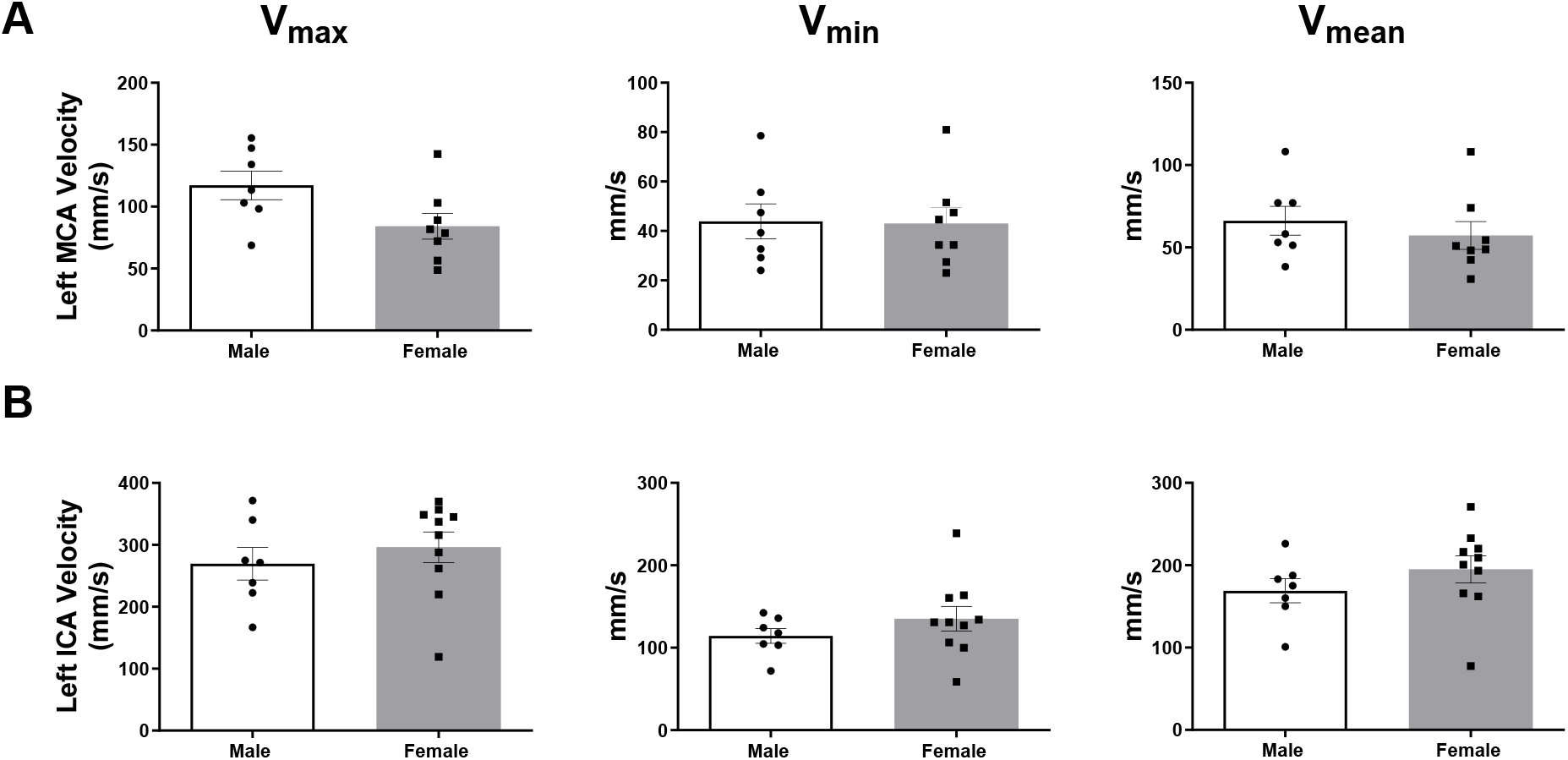
Cerebral Blood Flow Velocities of left MCA and ICA in adult Sprague-Dawley (SD) rats. Left MCA (**Panel A**) and ICA (**Panel B**) V_max_, V_min_, and V_mean_ in adult male versus female SD rats. Data are mean ± SEM, n=7-10.

### No Sex Differences in Systemic Arterial Stiffness in Adult SD Rats

There were no differences in thoracic or left common carotid artery PWV between female and male adult SD rats (**Figure 3A and B**).

**Figure 3.**
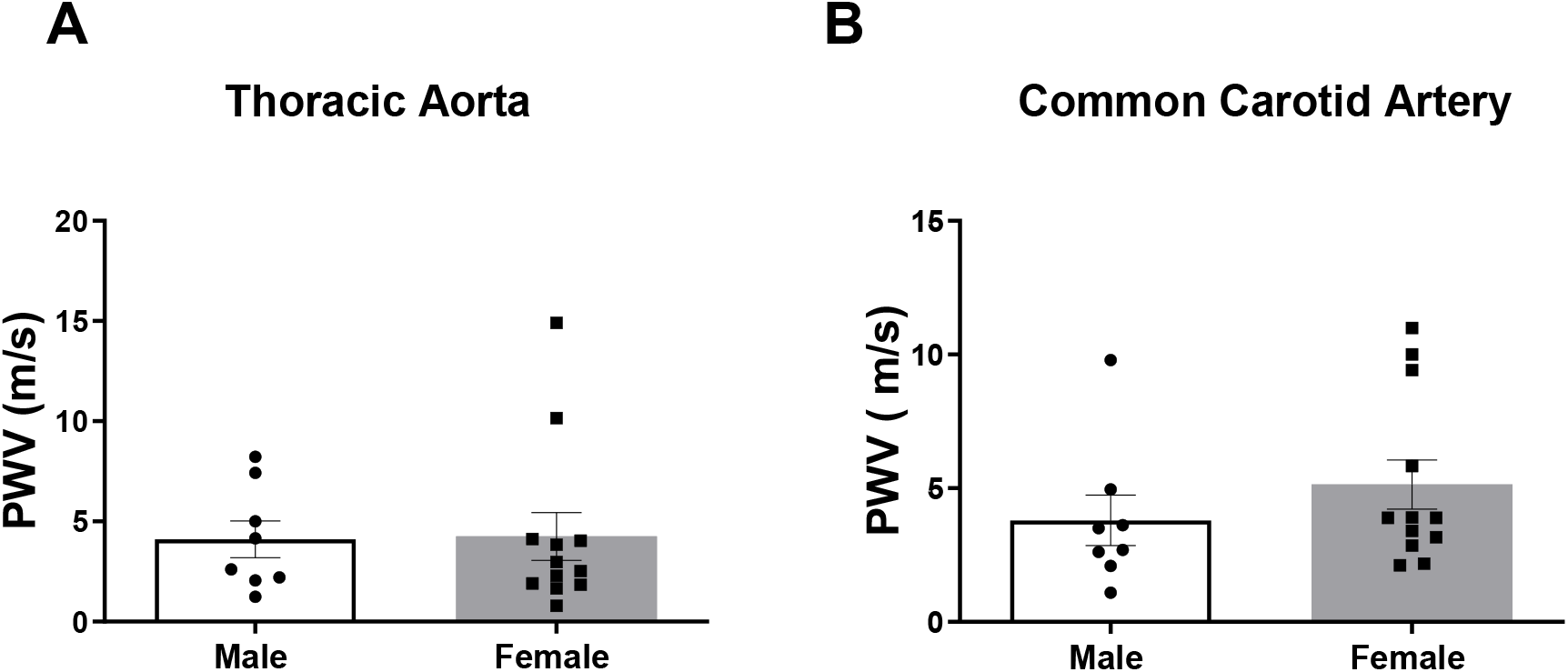
Aortic arch (**Panel A**) and left common carotid artery (**Panel B**) pulse wave velocity (PWV) in adult male versus female SD rats. Data are mean ± SEM, n=8-12.

### Sex Differences in the Levels of Vasoactive Targets in MCA

There were no differences in the levels of COX-2, PGIS, or TxA_2_-R in the endothelial cell layer of female versus male SD MCA (**Figures 4A, C, and D**). eNOS levels were elevated in the endothelial cell layer of LMCA of female versus male SD (**Figure 4B**). The levels TxA_2_-R were greater in the tunica media layer of female versus male SD (**Figure 4D**). There were no differences in the levels of COX-2 or PGIS in the tunica media of female versus male SD (**Figure 4A and C**).

**Figure 4.**
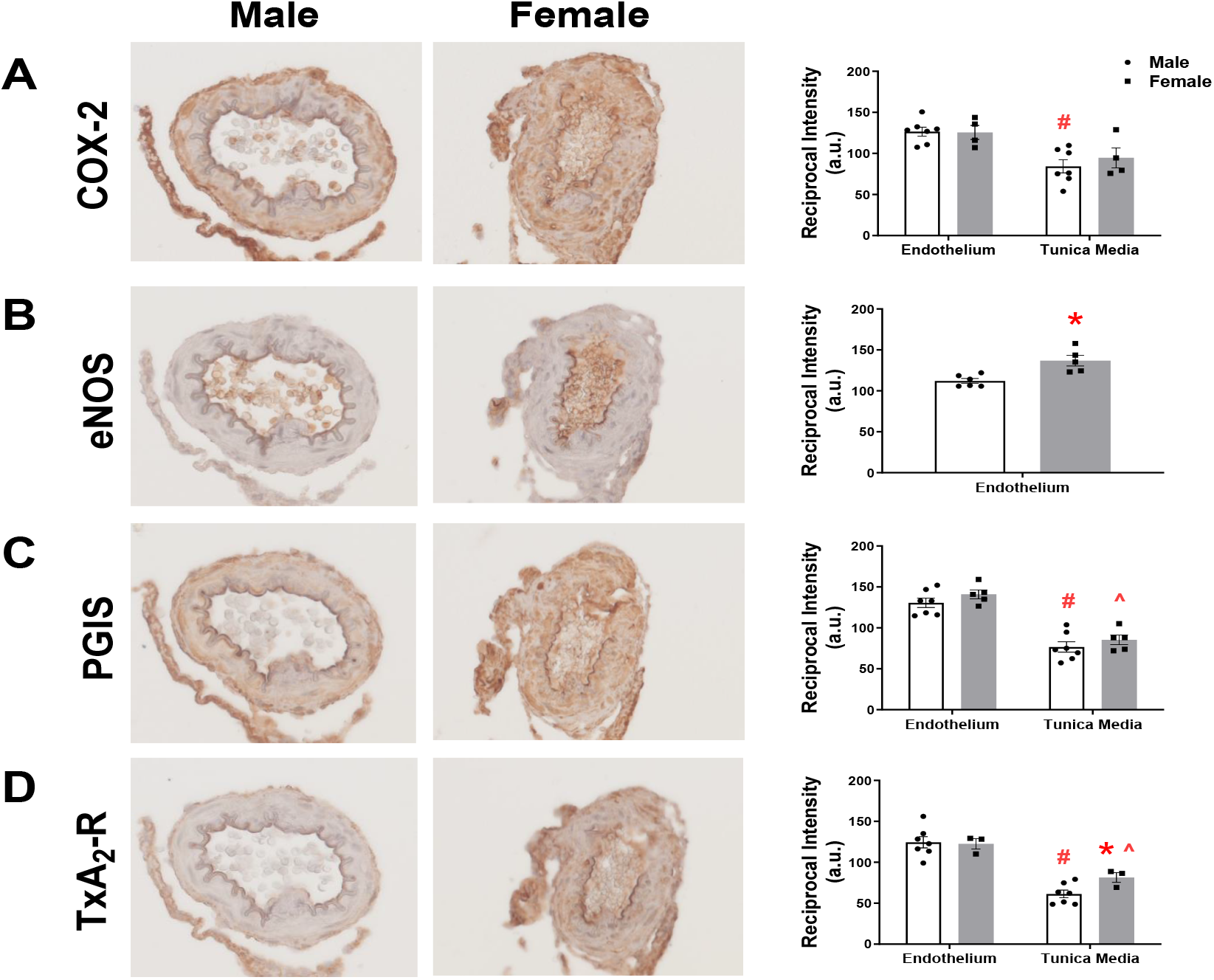
Representative images of immunohistochemical staining and the analysis of COX-2 (**Panel A**), eNOS (**Panel B**), PG synthase (**Panel C**) and TxA_2_-R (**Panel D**) in the endothelial and vascular smooth muscle cell layer of the left MCA in adult male versus female SD rats. Data are mean ± SEM; *p<0.05 vs. male tunica media, ^#^p<0.05 vs. male endothelium, ^p<0.05 vs. female endothelium; n=3-7.

### Sex Differences in Vascular Reactivity of Isolated MCA Segments

LMCAs from male and female SD rats showed similar optimal diameters and basal tone, with a lower active tone observed in female LMCA (**Table 2**). The vasodilatory response to acetylcholine was increased in female LMCA compared to male LMCA (**Figure 5; Table 3**). The response to the TxA_2_ agonist U-46619 was not different between male and female LMCA (**Figure 6A**), however in the presence of Indomethacin (10^−5^M) male LMCA showed a lower maximal response and sensitivity compared to female LMCA (**Figure 6B, Table 3**).

**Table 2.**
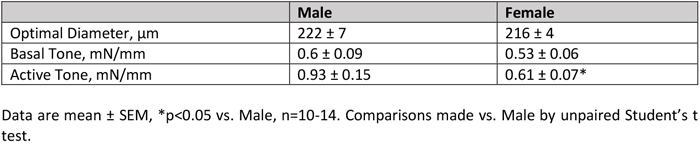
Structural and Functional Characteristics of LMCA in 25-Week-Old Male versus Female SD Rats.

**Table 3.**
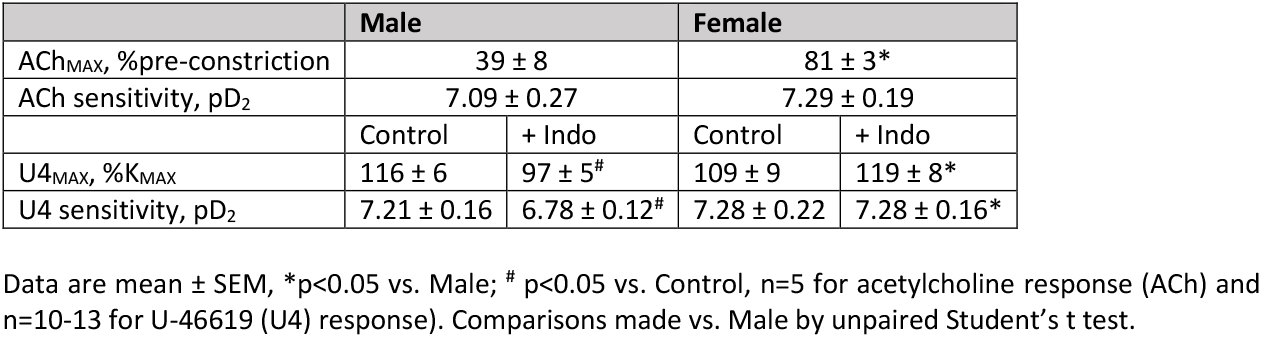
Responses to Acetylcholine (ACh) and U-46619 (U4) in LMCA in 25-Week-Old Male versus Female SD Rats.

**Figure 5.**
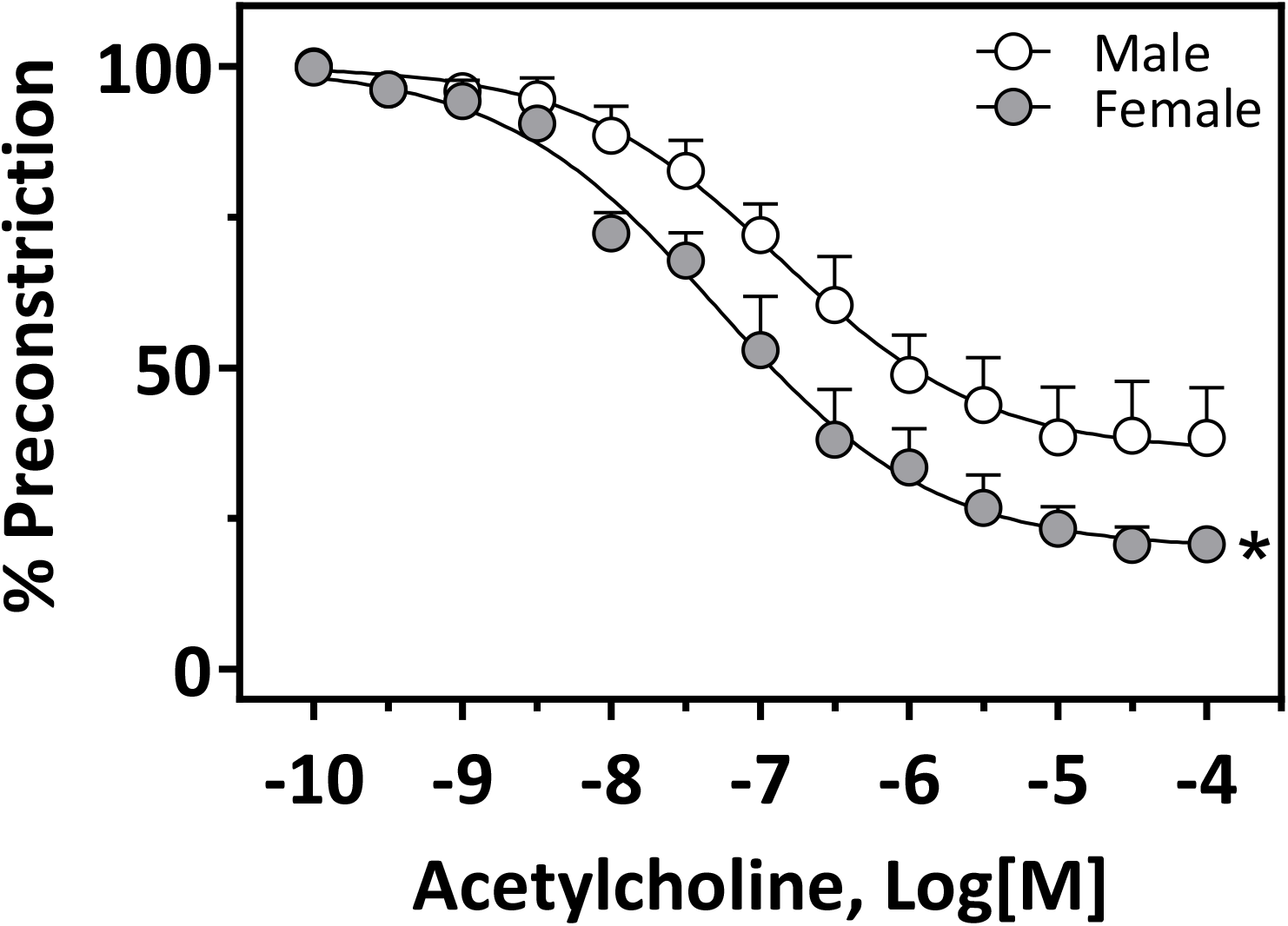
Vasodilatory response to acetylcholine in male and female LMCAs. LMCAs from male and female were pre-constricted with U-46619 (10^−7^M). *p<0.05 vs. male in maximal response; n=5 in each group.

**Figure 6.**
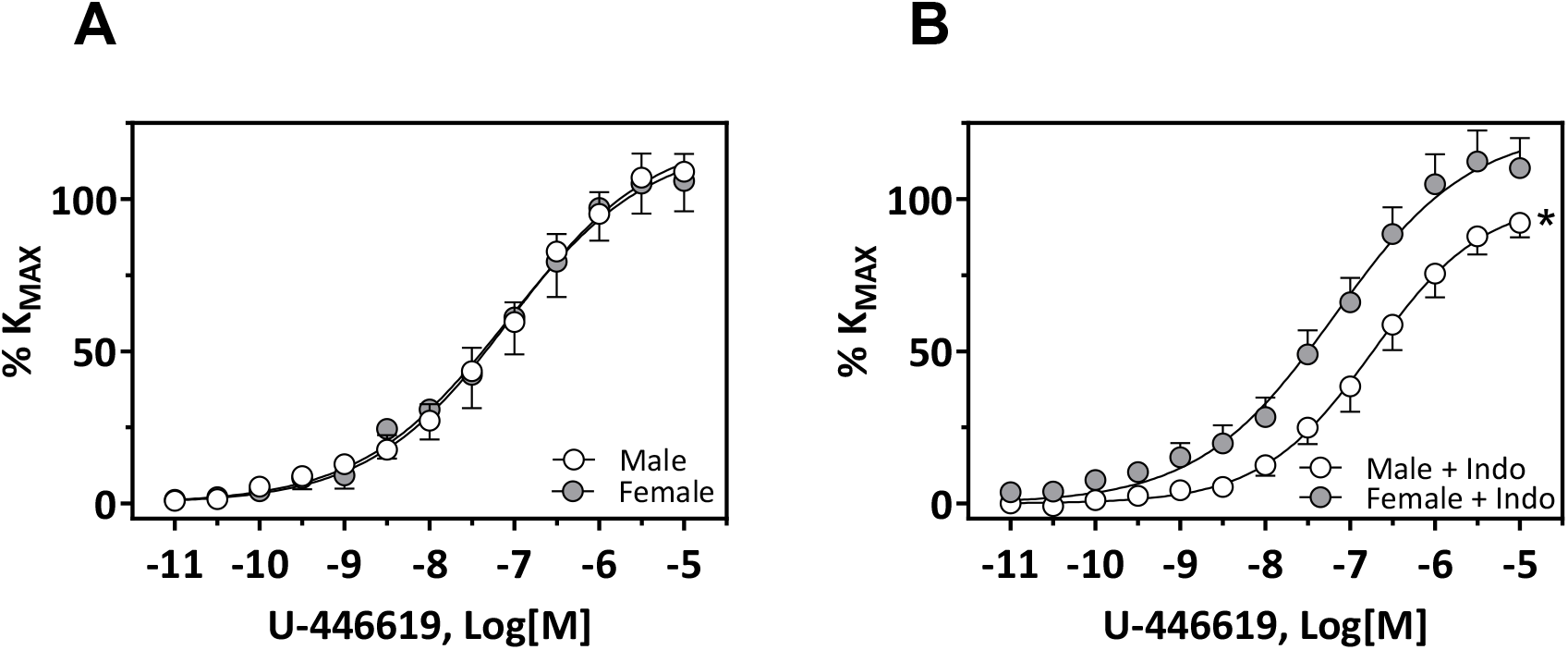
Contraction response to U-46619 in male and female LMCAs. LMCAs from male (n=10) and female (n=13) were exposed to U-46619 as indicated. Parallel experiments in intact arteries (A) and arteries preincubated with indomethacin (B; 10^−5^M) are shown. *p<0.05 vs. male in maximal response; n=10-13.

## DISCUSSION

Our study revealed that sex differences in MCA reactivity and hemodynamics in mature SD rats were not associated with changes in aortic or carotid PWV, a measure of systemic arterial stiffness, potentially indicating distinct regulatory mechanisms of cerebral versus systemic vascular resistance. Our data showed no differences in systolic or diastolic blood pressures but lower pulse pressure in female SD compared with males. Females had lower LMCA PI versus males, however there were no sex differences in thoracic aorta or common carotid artery PWV. Female LMCA had greater eNOS and TxA_2_R staining intensity in the endothelial and VSMC layers respectively, greater acetylcholine-dependent vasodilatation and TxA_2_R-dependent contraction compared with male LMCA.

In the present study, we used TCD ultrasound to evaluate LMCA resistance in 25-week-old SD rats. We focused on MCA because it is a major artery supplying blood to a significant portion of the brain. In addition, left MCA was more commonly pathologically affected in young adults (35). TCD ultrasound is a non-invasive method to evaluate cerebral artery structure and function. PI is considered a reliable marker of arterial resistance with higher PI indicating increased vascular resistance and hypoperfusion (23, 24, 36). Although there are few studies directly comparing changes in the PI in male versus female MCA, our findings are consistent with previous work demonstrating that young women have decreased MCA PI compared to age-matched men (37).

Pulsatile blood flow is often associated with end-organ damage (38). Large elastic arteries (including aorta and carotid arteries) serve to dampen pulsatile blood flow generated by the heart into a continuous flow. Given its need for steady flow and a relatively narrow range of acceptable perfusion pressures, the brain is particularly affected by the pulsatile blood flow. Stiffening of the large elastic arteries that occurs with age contributes to the increase in cerebral pulsatility that can potentially lead to cerebrovascular dysfunction and cognitive impairment (28). Pulse pressure and PWV increase with age and serve as predictors of cardiovascular disease including stroke and vascular dementia (12, 39). Compared with men, women experience accelerated vessel stiffening and pulsatile flow with aging. Our results are consistent with human studies where young women have lower PP compared to men (8, 13). Additionally, we found no sex difference in PWV of the thoracic aorta or common carotid artery consistent with previous studies evaluating abdominal aortic PWV in Dahl salt-resistant rats (40) or aortic arch PWV in glucose-fed or fructose- and high salt-treated SD rats (41). It is important to note that there are several different methods to calculate PWV. Similar to our study, Smulynan et al., found no sex differences in aortas of young men and women using the transit-time method, while Lefferts and colleagues used the local, one-point technique demonstrating reduced common carotid artery PWV in women compared with men prior to-but not after the menopause (8, 13). Thus, the vascular bed and the technique used to determine PWV need to be considered when making comparisons between different studies.

Regulation of cerebrovascular tone and blood flow is a complex process involving a number of mechanisms including: myogenic responses of smooth muscle to changes in arterial pressure, flow-metabolism coupling, autonomic nervous system input, vasoactive arachidonic acid (AA) metabolites, and endothelial factors such as eNOS and endothelium-derived hyperpolarizing factor (EDHF) (42-46). We show that female SD exhibit increased eNOS staining intensity in the endothelial layer of LMCA sections compared with males. Given that NO is a powerful vasodilator, sex differences in MCA resistance in our study are likely related to eNOS expression. Previous work demonstrates that estrogen has a dramatic influence on vascular function through modulation of eNOS (47). Human endothelial cells have intrinsic sexual dimorphism with females demonstrating higher eNOS expression and enzymatic activity (48). Estrogen increases eNOS enzymatic activity and levels in an estrogen-receptor dependent fashion (49-52). Additionally, PI in ICA increases after menopause while transdermal estrogen application reduces PI in ICA of postmenopausal women (53). It should be noted that we did not monitor the estrous cycle of rats which reflects the bioavailability of female sex hormones; however, the uterine weight-to-tibia length ratio was similar among female rats (0.1399 ± 0. 02 g/cm; 18.69% coefficient of variation). Other hormones, including testosterone and progesterone, also play important roles in vascular function. Progestins counteract the effect of estrogen on PI in carotid arteries (54). Chronic exposure to testosterone may promote cerebral artery vasoconstriction (19). In addition, the influence of sex hormones on other mechanisms (e.g. autonomic nervous system, flow-metabolism coupling) may also contribute to sex differences in the regulation of cerebrovascular function.

Many AA metabolites (e.g. prostaglandins, thromboxanes) are vasoactive with either vasoconstrictor or vasodilator effects (55). Isoforms of the cyclooxygenase (COX) enzyme are primarily responsible for the initial metabolism of AA; subsequent processing via thromboxane synthase or PGIS produces vasoactive metabolites including thromboxane A_2_ (TxA_2_) or prostacyclin (PGI_2_). We found no significant difference between sexes in the expression of COX-2. Additionally, there were no differences between sexes in the expression of PGIS, the enzyme responsible for producing the powerful vasodilator PGI_2_. However, the levels of TxA_2_R were increased in the VSMC layer of female SD. Considering that female rats exhibit less VSMCs in the MCA, it is possible the total number of receptors is similar between male and female SD (7). Although, we also did not measure the levels of thromboxane synthase, the enzyme responsible for producing TxA_2,_ it is possible that the overall availability of TxA_2_ may differ between sexes due to differential expression of thromboxane synthase. In contrast to endothelial cells, increased TxA_2_R levels in VSMC of females MCA does not appear to influence COX-2 expression. Although the products of AA metabolism are thought to play a somewhat minor role in maintaining cerebrovascular tone (42, 44, 45), the physiologic significance of increased TxA_2_R in the VSMC layer of female MCA in relation to female sex hormone status needs further investigation (56).

Our data also showed differences in local vascular responses between isolated male and female LMCAs. The lower active tone in female LMCAs may reflect their previously reported lower smooth muscle content (7), whereas the increased expression of eNOS may explain the increased vasodilatory response we observed in female LMCAs in our study. This increased vasodilatory response may counteract increased levels of the TXA_2_ receptor and be responsible for the similar responses to U-46619 we observed in male and female LMCAs. Pre-incubation with the unspecific blocker of the prostanoid pathway indomethacin should decrease the levels of all vasoactive products of AA metabolism, and in this scenario the increased levels of the TXA_2_ receptor in female LMCAs would also be responsible for the lower U-46619-dependent contraction in male but not in female LMCAs. Overall, we demonstrated sex differences in MCA reactivity and hemodynamics independent from changes in systemic arterial stiffening in middle-aged SD rats potentially indicating diverse mechanisms underlying vascular resistance in cerebral versus systemic arterial beds.

## DATA AVAILABILITY

The data are available from the corresponding authors upon reasonable request.

## ACKNOWLEDGMENTS

We acknowledge the Preclinical Ultrasound and Photoacoustic Imaging Core of Wake Forest University School of Medicine.

## DISCLAIMERS

None

## AUTHOR CONTRIBUTIONS

JWR conceived and designed research, performed experiments, analyzed data, interpreted results of experiments, prepared figures, drafted manuscript, and edited and revised manuscript. XS performed experiments, analyzed data, interpreted results of experiments, edited and revised manuscript. NCD performed experiments, analyzed data, interpreted results of experiments. VMP conceived and designed research, performed experiments, analyzed data, interpreted results of experiments, prepared figures, drafted manuscript, and edited and revised manuscript, and approved final version of manuscript. LMY conceived and designed research, analyzed data, interpreted results of experiments, prepared figures, edited and revised manuscript, and approved final version of manuscript.

## REFERENCES

1. Bushnell C, McCullough LD, Awad IA, Chireau MV, Fedder WN, Furie KL, Howard VJ, Lichtman JH, Lisabeth LD, Pina IL, Reeves MJ, Rexrode KM, Saposnik G, Singh V, Towfighi A, Vaccarino V, Walters MR, American Heart Association Stroke C, Council on C, Stroke N, Council on Clinical C, Council on E, Prevention, and Council for High Blood Pressure R. Guidelines for the prevention of stroke in women: a statement for healthcare professionals from the American Heart Association/American Stroke Association. Stroke 45: 1545–1588, 2014.

2. Petrea RE, Beiser AS, Seshadri S, Kelly-Hayes M, Kase CS, and Wolf PA. Gender differences in stroke incidence and poststroke disability in the Framingham heart study. Stroke 40: 1032–1037, 2009.

3. Seshadri S, Beiser A, Kelly-Hayes M, Kase CS, Au R, Kannel WB, and Wolf PA. The lifetime risk of stroke: estimates from the Framingham Study. Stroke 37: 345–350, 2006.

4. Gao Sh, H.C.; Hall, K.S.; Hui, S. The Relationships Between Age, Sex, and the Incidence of Dementia and Alzheimer Disease Arch Gen Psychiatry 55: 809–815, 1998.

5. Read Sp, N.L.; Gatz, M.; Berg, S.; Vuoksimaa, E.; Malmberg, B.; Johansson, B.; McClearn, G.E. Sex Differences After All Those Years? Heritability of Cognitive Abilities in Old Age Journal of Gerontology 61B: 137–143, 2006.

6. Navarro-Orozco D, and Sanchez-Manso JC. Neuroanatomy, Middle Cerebral Artery. In: StatPearls. StatPearls [Internet]: Treasure Island (FL): StatPearls Publishing, 2024.

7. Wang S, Zhang H, Liu Y, Li L, Guo Y, Jiao F, Fang X, Jefferson JR, Li M, Gao W, Gonzalez-Fernandez E, Maranon RO, Pabbidi MR, Liu R, Alexander BT, Roman RJ, and Fan F. Sex differences in the structure and function of rat middle cerebral arteries. Am J Physiol Heart Circ Physiol 318: H1219–H1232, 2020.

8. Lefferts WK, DeBlois JP, Augustine JA, Keller AP, and Heffernan KS. Age, sex, and the vascular contributors to cerebral pulsatility and pulsatile damping. J Appl Physiol (1985) 129: 1092–1101, 2020.

9. de la Cruz-Cosme C, Dawid-Milner MS, Ojeda-Burgos G, Gallardo-Tur A, and Segura T. Doppler Resistivity and Cerebral Small Vessel Disease: Hemodynamic Structural Correlation and Usefulness for the Etiological Classification of Acute Ischemic Stroke. J Stroke Cerebrovasc Dis 27: 3425–3435, 2018.

10. Lau KK, Pego P, Mazzucco S, Li L, Howard DP, Kuker W, and Rothwell PM. Age and sex-specific associations of carotid pulsatility with small vessel disease burden in transient ischemic attack and ischemic stroke. Int J Stroke 13: 832–839, 2018.

11. Chung CP, Lee HY, Lin PC, and Wang PN. Cerebral Artery Pulsatility is Associated with Cognitive Impairment and Predicts Dementia in Individuals with Subjective Memory Decline or Mild Cognitive Impairment. J Alzheimers Dis 60: 625–632, 2017.

12. Mitchell GF, Gudnason V, Launer LJ, Aspelund T, and Harris TB. Hemodynamics of increased pulse pressure in older women in the community-based Age, Gene/Environment Susceptibility-Reykjavik Study. Hypertension 51: 1123–1128, 2008.

13. Smulyan H, Asmar RG, Rudnicki A, London GM, and Safar ME. Comparative effects of aging in men and women on the properties of the arterial tree. J Am Coll Cardiol 37: 1374–1380, 2001.

14. Tarumi T, Ayaz Khan M, Liu J, Tseng BY, Parker R, Riley J, Tinajero C, and Zhang R. Cerebral hemodynamics in normal aging: central artery stiffness, wave reflection, and pressure pulsatility. J Cereb Blood Flow Metab 34: 971–978, 2014.

15. Yang D, Cabral D, Gaspard EN, Lipton RB, Rundek T, and Derby CA. Cerebral Hemodynamics in the Elderly: A Transcranial Doppler Study in the Einstein Aging Study Cohort. J Ultrasound Med 35: 1907–1914, 2016.

16. Purkayastha S, Fadar O, Mehregan A, Salat DH, Moscufo N, Meier DS, Guttmann CR, Fisher ND, Lipsitz LA, and Sorond FA. Impaired cerebrovascular hemodynamics are associated with cerebral white matter damage. J Cereb Blood Flow Metab 34: 228–234, 2014.

17. Wahlin A, Ambarki K, Birgander R, Malm J, and Eklund A. Intracranial pulsatility is associated with regional brain volume in elderly individuals. Neurobiol Aging 35: 365–372, 2014.

18. Lisabeth L, and Bushnell C. Stroke risk in women: the role of menopause and hormone therapy. Lancet Neurol 11: 82–91, 2012.

19. Robison LS, Gannon OJ, Salinero AE, and Zuloaga KL. Contributions of sex to cerebrovascular function and pathology. Brain Res 1710: 43–60, 2019.

20. Shekhar S, Travis OK, He X, Roman RJ, and Fan F. Menopause and Ischemic Stroke: A Brief Review. MOJ Toxicol 3: 2017.

21. Girijala RL, Sohrabji F, and Bush RL. Sex differences in stroke: Review of current knowledge and evidence. Vasc Med 22: 135–145, 2017.

22. Roy-O’Reilly M, and McCullough LD. Age and Sex Are Critical Factors in Ischemic Stroke Pathology. Endocrinology 159: 3120–3131, 2018.

23. Kassab MY, Majid A, Farooq MU, Azhary H, Hershey LA, Bednarczyk EM, Graybeal DF, and Johnson MD. Transcranial Doppler: an introduction for primary care physicians. J Am Board Fam Med 20: 65–71, 2007.

24. Wielicka MN-G, J.; Kozera, G.; Bieniaszewski, L. Clinical application of pulsatility index. Med Res J 5: 201–210, 2020.

25. Giustetto P, Filippi M, Castano M, and Terreno E. Non-invasive parenchymal, vascular and metabolic high-frequency ultrasound and photoacoustic rat deep brain imaging. J Vis Exp 2015.

26. Hussein AE, Brunozzi D, Shakur SF, Ismail R, Charbel FT, and Alaraj A. Cerebral Aneurysm Size and Distal Intracranial Hemodynamics: An Assessment of Flow and Pulsatility Index Using Quantitative Magnetic Resonance Angiography. Neurosurgery 83: 660–665, 2018.

27. Harris S, Reyhan T, Ramli Y, Prihartono J, and Kurniawan M. Middle Cerebral Artery Pulsatility Index as Predictor of Cognitive Impairment in Hypertensive Patients. Front Neurol 9: 538, 2018.

28. Reeve EH, Barnes JN, Moir ME, and Walker AE. Impact of arterial stiffness on cerebrovascular function: a review of evidence from humans and preclincal models. Am J Physiol Heart Circ Physiol 326: H689–H704, 2024.

29. Williams R, Needles A, Cherin E, Zhou YQ, Henkelman RM, Adamson SL, and Foster FS. Noninvasive ultrasonic measurement of regional and local pulse-wave velocity in mice. Ultrasound Med Biol 33: 1368–1375, 2007.

30. Elsangeedy E, Yamaleyeva DN, Edenhoffer NP, Deak A, Soloshenko A, Ray J, Sun X, Shaltout OH, Diaz NC, Westwood B, Kim-Shapiro D, Diz DI, Soker S, Pulgar VM, Ronca A, Willey JS, and Yamaleyeva LM. Sex-Specific Cardiovascular Adaptations to Simulated Microgravity in Sprague-Dawley Rats. bioRxiv 2024.2003.2029.587264, 2024.

31. Pulgar VM, Yamaleyeva LM, Varagic J, McGee C, Bader M, Dechend R, and Brosnihan KB. Functional changes in the uterine artery precede the hypertensive phenotype in a transgenic model of hypertensive pregnancy. Am J Physiol Endocrinol Metab 309: E811–817, 2015.

32. Pulgar VM, Yamaleyeva LM, Varagic J, McGee CM, Bader M, Dechend R, Howlett AC, and Brosnihan KB. Increased angiotensin II contraction of the uterine artery at early gestation in a transgenic model of hypertensive pregnancy is reduced by inhibition of endocannabinoid hydrolysis. Hypertension 64: 619–625, 2014.

33. Nguyen DHZ, T.; Shu, J.; Mao, J. Quantifying chromogen intensity in immunohistochemistry via reciprocal intensity. Cancer InCytes 2: 2013.

34. Pulgar VM, Yasuda M, Gan L, Desnick RJ, and Bonkovsky HL. Sex differences in vascular reactivity in mesenteric arteries from a mouse model of acute intermittent porphyria. Molecular Genetics and Metabolism 128: 376–381, 2019.

35. Naess H, Waje-Andreassen U, Thomassen L, and Myhr K-M. High Incidence of Infarction in the Left Cerebral Hemisphere Among Young Adults. Journal of Stroke and Cerebrovascular Diseases 15: 241–244, 2006.

36. Cho Sjs, Y.H.; Kim, G.W.; Kim, J. Blood flow velocity changes in the middle cerebral artery as an index of the chronicity of hypertension. J Neurol Sci 150: 77–80, 1997.

37. Alwatban MR, Aaron SE, Kaufman CS, Barnes JN, Brassard P, Ward JL, Miller KB, Howery AJ, Labrecque L, and Billinger SA. Effects of age and sex on middle cerebral artery blood velocity and flow pulsatility index across the adult lifespan. J Appl Physiol (1985) 130: 1675–1683, 2021.

38. Mitchell GF. Aortic stiffness, pressure and flow pulsatility, and target organ damage. J Appl Physiol (1985) 125: 1871–1880, 2018.

39. Waldstein SR, Rice SC, Thayer JF, Najjar SS, Scuteri A, and Zonderman AB. Pulse pressure and pulse wave velocity are related to cognitive decline in the Baltimore Longitudinal Study of Aging. Hypertension 51: 99–104, 2008.

40. Decano JL, Pasion KA, Black N, Giordano NJ, Herrera VL, and Ruiz-Opazo N. Sex-specific genetic determinants for arterial stiffness in Dahl salt-sensitive hypertensive rats. BMC Genet 17: 19, 2016.

41. Komnenov D, and Rossi NF. Fructose-induced salt-sensitive blood pressure differentially affects sympathetically mediated aortic stiffness in male and female Sprague-Dawley rats. Physiol Rep 11: e15687, 2023.

42. Cipolla MJ. The Cerebral Circulation. San Rafael, CA: Morgan & Claypool Life Sciences, 2009.

43. Peterson EC, Wang Z, and Britz G. Regulation of cerebral blood flow. Int J Vasc Med 2011: 823525, 2011.

44. You J, Golding EM, and Bryan RM, Jr. Arachidonic acid metabolites, hydrogen peroxide, and EDHF in cerebral arteries. Am J Physiol Heart Circ Physiol 289: H1077–1083, 2005.

45. Andresen J, Shafi NI, and Bryan RM, Jr. Endothelial influences on cerebrovascular tone. J Appl Physiol (1985) 100: 318–327, 2006.

46. Koep JL, Taylor CE, Coombes JS, Bond B, Ainslie PN, and Bailey TG. Autonomic control of cerebral blood flow: fundamental comparisons between peripheral and cerebrovascular circulations in humans. J Physiol 600: 15–39, 2022.

47. Chambliss KL, and Shaul PW. Estrogen modulation of endothelial nitric oxide synthase. Endocr Rev 23: 665–686, 2002.

48. Cattaneo MG, Vanetti C, Decimo I, Di Chio M, Martano G, Garrone G, Bifari F, and Vicentini LM. Sex-specific eNOS activity and function in human endothelial cells. Sci Rep 7: 9612, 2017.

49. MacRitchie AN, Jun SS, Chen Z, German Z, Yuhanna IS, Sherman TS, and Shaul PW. Estrogen upregulates endothelial nitric oxide synthase gene expression in fetal pulmonary artery endothelium. Circ Res 81: 355–362, 1997.

50. Hayashi TY, K.; Esaki, T.; Kuzuya, M.; Satake, S.; Ishikawa, T.; Hidaka, H.; Iguchi, A. Estrogen increases endothelial nitric oxide by a receptor-mediated system. Biochem Biohphys Res Commun 214: 847–855, 1995.

51. Sumi DH, T.; Jayachandran, M.; Iguchi, A. Estrogen prevents destabilization of endothelial nitric oxide synthase mRNA induced by tumor necrosis factor α through estrogen receptor mediated system. Life Sci 69: 1651–1660, 2001.

52. Sumi Di, L.J. Estrogen-related receptor α1 up-regulates endothelial nitric oxide synthase expression Proc Natl Acad Sci U S A 100: 14451–14456, 2003.

53. Gangar KF, Vyas S, Whitehead M, Crook D, Meire H, and Campbell S. Pulsatility index in internal carotid artery in relation to transdermal oestradiol and time since menopause. Lancet 338: 839–842, 1991.

54. Luckas Mjmg, T.; Biljan, M.M.; Buckett, W.M.; Aird, I.A.; Drakeley, A.; Kingsland, C.R. The effect of progestagens on the carotid artery pulsatility index in postmenopausal women on oestrogen replacement therapy. Eur J Obstet Gynecol Reprod Biol 76: 221–224, 1998

55. Bogatcheva NV, Sergeeva MG, Dudek SM, and Verin AD. Arachidonic acid cascade in endothelial pathobiology. Microvasc Res 69: 107–127, 2005.

56. Ospina JA, Duckles SP, and Krause DN. 17beta-estradiol decreases vascular tone in cerebral arteries by shifting COX-dependent vasoconstriction to vasodilation. Am J Physiol Heart Circ Physiol 285: H241–250, 2003.

